# Chimpanzee Brain Morphometry Utilizing Standardized MRI Preprocessing and Macroanatomical Annotations

**DOI:** 10.1101/2020.04.20.046680

**Authors:** Sam Vickery, William D. Hopkins, Chet C. Sherwood, Steven J. Schapiro, Robert D. Latzman, Svenja Caspers, Christian Gaser, Simon B. Eickhoff, Robert Dahnke, Felix Hoffstaedter

## Abstract

Chimpanzees are among the closest living relatives to humans and, as such, provide a crucial comparative model for investigating primate brain evolution. In recent years, human brain mapping has strongly benefited from enhanced computational models and image processing pipelines that could also improve data analyses in animals by using species-specific templates. In this study, we use structural MRI data from the National Chimpanzee Brain Resource (NCBR) to develop the chimpanzee brain reference template Juna.Chimp for spatial registration and the macro-anatomical brain parcellation Davi130 for standardized whole-brain analysis. Additionally, we introduce a ready-to-use image processing pipeline built upon the CAT12 toolbox in SPM12, implementing a standard human image preprocessing framework in chimpanzees. Applying this approach to data from 178 subjects, we find strong evidence for age-related GM atrophy in multiple regions of the chimpanzee brain, as well as, a human-like anterior-posterior pattern of hemispheric asymmetry in medial chimpanzee brain regions.

## Introduction

Chimpanzees (*Pan troglodytes*) along with bonobos (*Pan paniscus*) represent the closest extant relatives of humans sharing a common ancestor approximately 7-8 million years ago (Langergraber et al. 2012). Experimental and observational studies, in both the field and in captivity, have documented a range of cognitive abilities that are shared with humans such as tool use and manufacturing (Shumaker et al. 2011), symbolic thought (de Waal 1996), mirror self-recognition (Anderson and Gallup 2015; Hecht et al. 2017) and some basic elements of language (Savage-Rumbaugh 1986; Savage-Rumbaugh and Lewin 1994; Tomasello and Call 1997) like conceptual metaphorical mapping (Dahl and Adachi 2013). This cognitive complexity together with similar neuroanatomical features (Zilles et al. 1989; Rilling and Insel 1999; Gomez-Robles et al. 2013; Hopkins et al. 2014a, 2017) and genetic proximity (Waterson et al. 2005) renders these species unique among non-human primates to study the evolutional origins of the human condition. In view of evolutionary neurobiology, the relatively recent divergence between humans and chimpanzees explains the striking similarities in major gyri and sulci, despite profound differences in overall brain size. Numerous studies using magnetic resonance imaging (MRI) have compared relative brain size, shape, and gyrification in humans and chimpanzees (Zilles et al. 1989; Rilling and Insel 1999; Gomez-Robles et al. 2013; Hopkins et al. 2014b, 2017).

Previous studies of brain aging in chimpanzees have reported minimal indications of atrophy (Herndon et al. 1999; Sherwood et al. 2011; Chen et al. 2013; Autrey et al. 2014). Nevertheless, Edler and colleagues (2017) recently found that brains of older chimpanzees’ exhibit both neurofibrillary tangles and amyloid plaques, the classical features of Alzheimer’s disease (AD). Neurodegeneration in the aging human brain includes marked atrophy in frontal and temporal lobes and decline in glucose metabolism even in the absence of detectable amyloid beta deposition, which increases the likelihood of cognitive decline and development of AD (Jagust 2018). Given the strong association of brain atrophy and amyloid beta in humans, this phenomenon requires further investigation in chimpanzees.

Cortical asymmetry is a prominent feature of brain organization in many primate species (Hopkins et al. 2015) and was recently shown in humans in a large scale ENIGMA (Enhancing Neuroimaging Genetics through Meta-Analysis) study (Kong et al. 2018). For chimpanzees, various studies have reported population-level asymmetries in different parts of the brain associated with higher order cognitive functions like tool-use (Freeman et al. 2004; Hopkins et al. 2008, 2017; Hopkins and Nir 2010; Lyn et al. 2011; Bogart et al. 2012; Gilissen and Hopkins 2013) but these results are difficult to compare within and across species, due to the lack of standardized registration and parcellation techniques as found for humans.

To date, there is no common reference space for the chimpanzee brain available to reliably associate and quantitatively compare neuro-anatomical evidence, nor is there a standardized image processing protocol for T1-weighted brain images from chimpanzees that matches human imaging standards. With the introduction of voxel-based morphometry (VBM, Ashburner and Friston 2000) and the ICBM (international consortium of brain mapping) standard human reference brain templates almost two decades ago (Mazziotta et al. 2001) MRI analyses became directly comparable and generally reproducible In this study we adapt state-of-the-art MRI (magnetic resonance imaging) processing methods to assess brain aging and cortical asymmetry in the chimpanzee brain. To make this possible, we rely on the largest openly available resource of chimpanzee MRI data: the *National Chimpanzee Brain Resource* (NCBR, www.chimpanzeebrain.org), including in vivo MRI images of 223 subjects from 9 to 54 years of age (Mean age = 26.9 ± 10.2 years). The aim of this study is the creation of a chimpanzee template permitting automated and reproducible image registration, normalization, statistical analysis and visualization to systematically investigate brain aging and hemispheric asymmetry in chimpanzees.

## Results

Initially, we created the population based Juna (Forschungszentrum <u>Jue</u>lich - University Je<u>na</u>) T1-template, tissue probability maps (TPM) for tissue classification and a non-linear spatial registration ‘Shooting’ templates (Figure 1) in an iterative fashion at 1mm spatial resolution. The preprocessing pipeline and templates creation were established using the freely available *Statistical Parametric Mapping* (SPM12 v7487, http://www.fil.ion.ucl.ac.uk/spm/) software and *Computational Anatomy* Toolbox (CAT12 r1434, http://www.neuro.uni-jena.de/cat/). Juna.Chimp templates and the Davi130 parcellation as well as images for analysis are interactively accessible via the Juna.Chimp web viewer (http://junachimp.inm7.de/).

**Figure 1.**
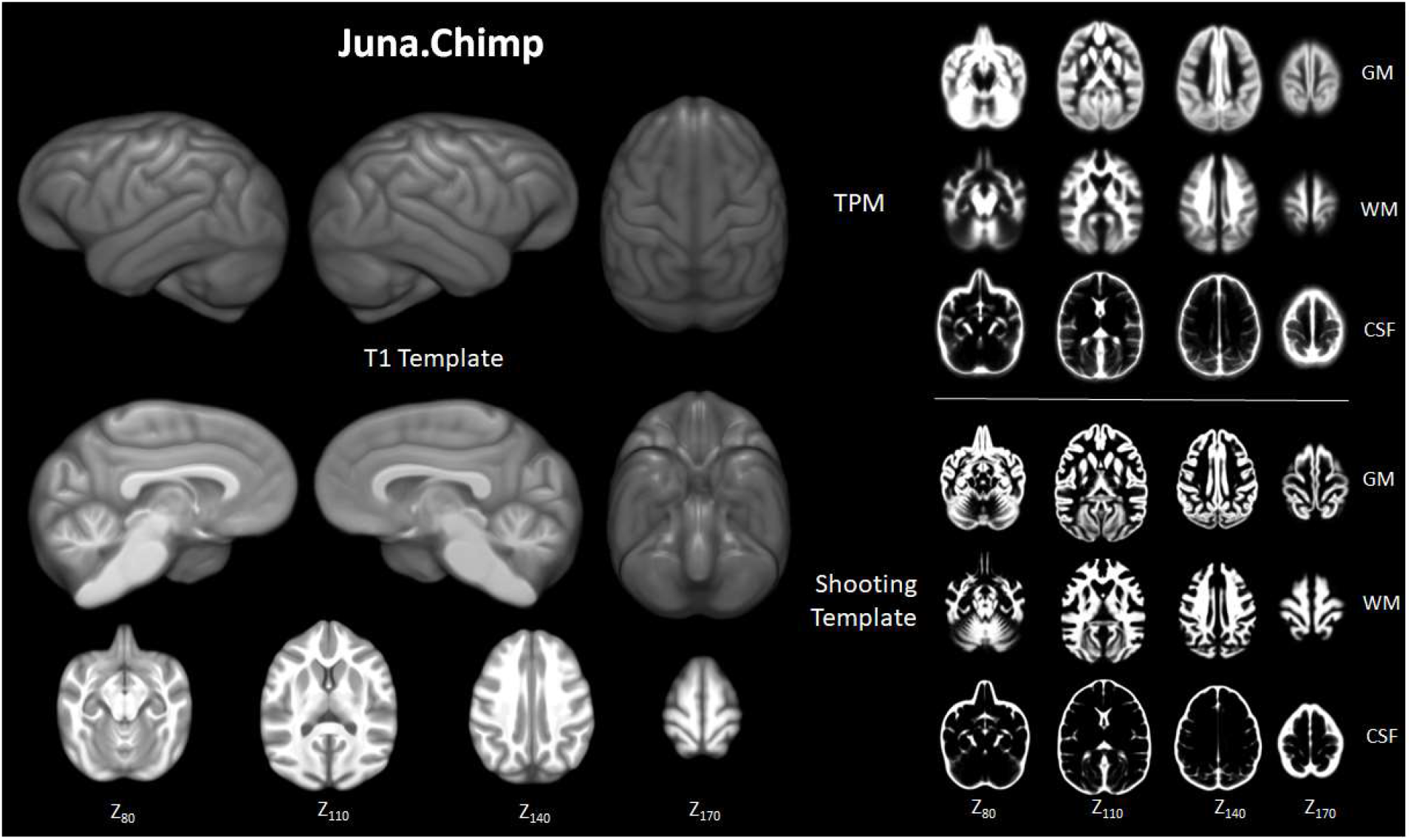
Juna.Chimp templates including the average T1-template, tissue probability maps (TPM), and Geodesic Shooting template. For Shooting templates and TPM axial slices are shown of gray matter (GM), white matter (WM), and cerebrospinal fluid (CSF). All templates are presented at 0.5mm resolution.

To enable more direct comparison to previous research, we manually created the Davi130 parcellation (by R.D. and S.V.), a whole brain macroanatomical annotation based on the Juna T1 template (Figure 2). The delineation of regions within the cortex was determined by following major gyri and sulci, whereby, large regions were arbitrarily split into two to three sub-regions of approximate equal size even though histological studies show that micro-anatomical borders between brain regions are rarely situated at the fundus (Sherwood et al. 2003; Schenker et al. 2010; Spocter et al. 2010; Amunts and Zilles 2015)‥ This process yielded 65 regions per hemisphere for a total of 130 regions for the Davi130 macro-anatomical manual parcellation (Figure 2, Supplementary Table 1).

**Figure 2.**
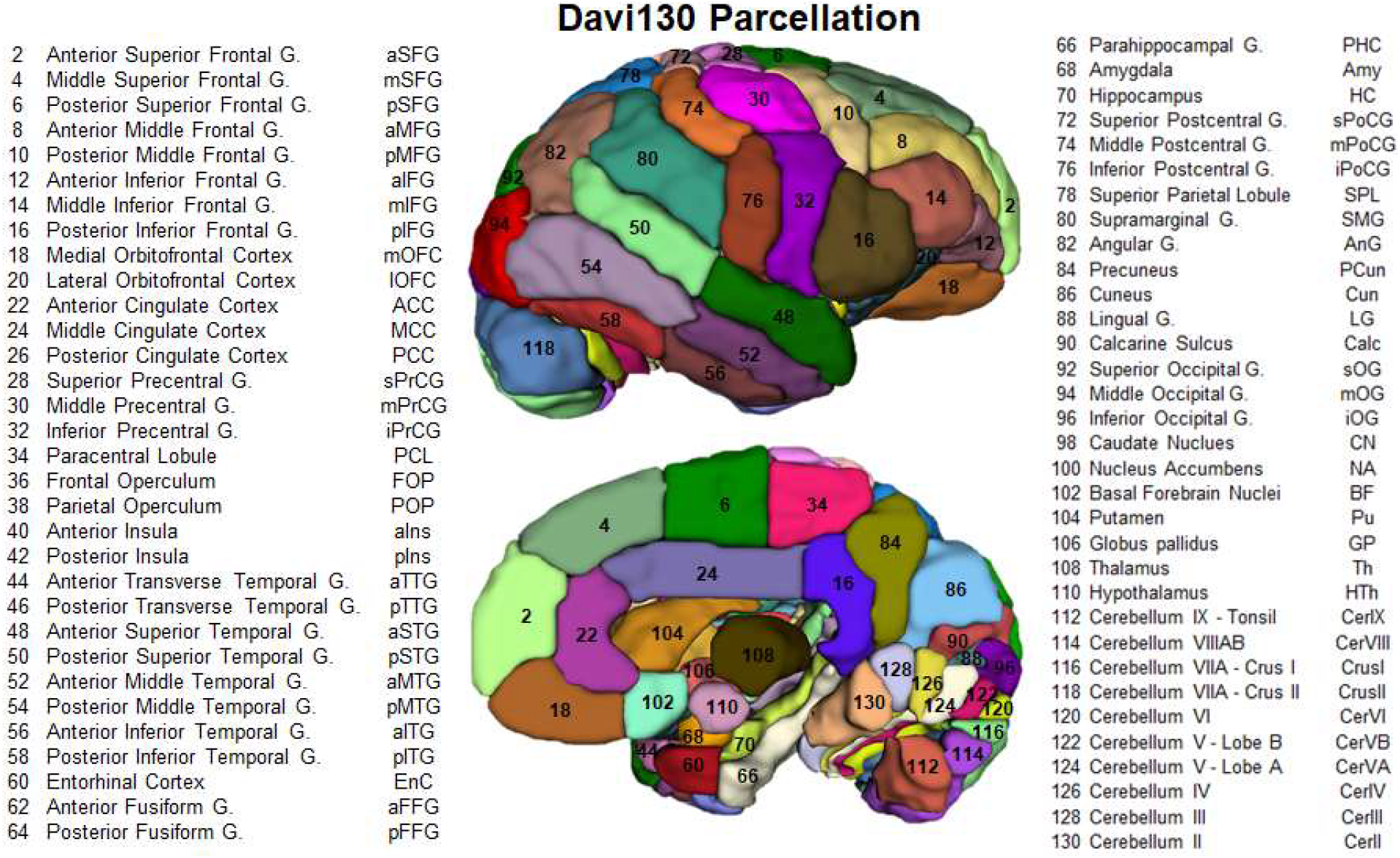
Lateral and medial aspect of the Davi130 parcellation right hemisphere. Visible regions are numbered with Davi130 parcellation region numbers and correspond to names in the figure. Odd numbers correspond to left hemisphere regions while even numbers are in the right hemisphere.

Following successful CAT12 preprocessing rigorous quality control (QC) was employed to identify individual MRI scans suitable for statistical analysis of brain aging and hemispheric asymmetry in chimpanzees. Our final sample consists of 178 chimpanzees including 120 females with an age range of 11 to 54 years and a mean age of 26.7 ± 9.8 years (Figure 3A). Correlation analysis between GM fraction of total intracranial volume and age revealed a significant negative association between the two (R^2^ = 0.11, p < 0.0001) demonstrating age-related decline in overall GM (Figure 3B). Both male and female subjects show a significant age effect on GM (male: R^2^ = 0.08, p = 0.03; female R^2^ = 0.11, p = 0.0001). The linear model showed no significant sex differences of GM decline (p = 0.08).

**Figure 3.**
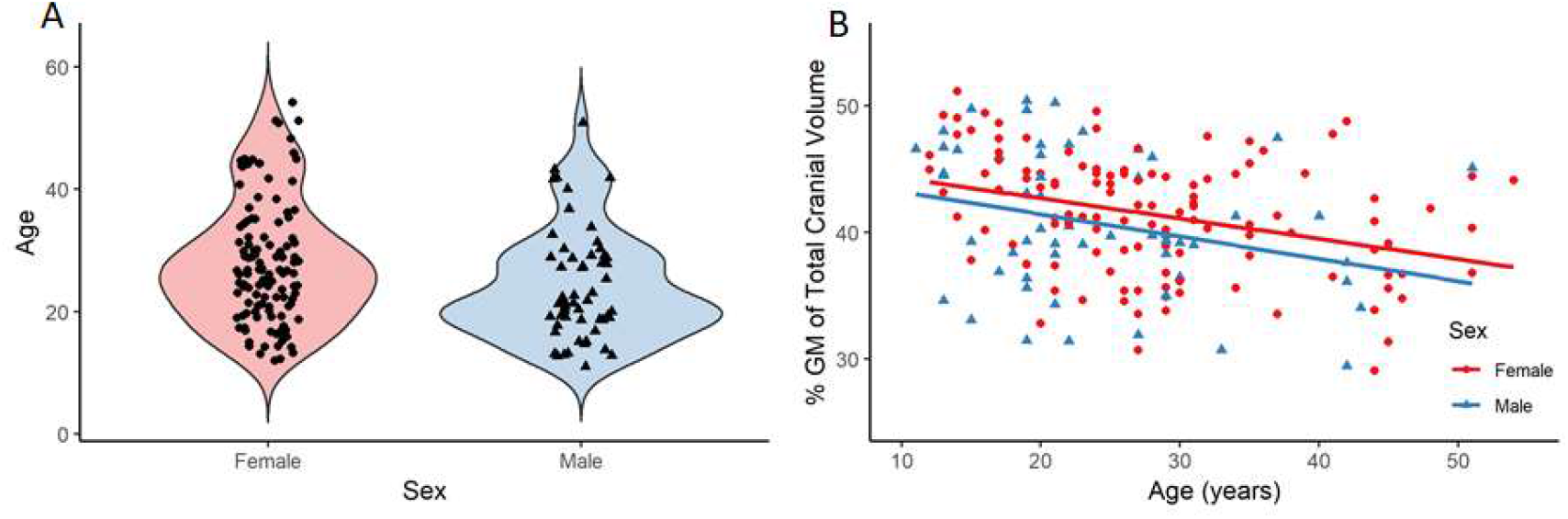
A - Distribution of age and sex in the final sample of 178 chimpanzees. B-linear relationship between GM and age for female and male respectively.

Region based morphometry analysis was applied to test for local effect of age on GM. Linear regression analyses identified 23 regions in the Davi130 parcellation across both hemispheres that were significantly associated with age after family-wise error (FWE) correction for multiple testing (Figure 4, for all region results see Supplementary). Specifically, GM decline with age was found bilaterally in the lateral orbitofrontal cortex (lOFC) and mid-cingulate (MCC) as well as unilaterally in the right precuneus (PCun), posterior cingulate (PCC), and lingual gyrus (LG), in addition to the left middle superior frontal gyrus(mSFG), anterior transverse temporal gyrus (aTTG) and calcarine sulcus (Calc) within the cerebral cortex. The strongest association with aging was in the bilateral putamen (Pu) and caudate nucleus (CN), while the nucleus accumbens (NA) and superior cerebellum (Ce-rIII, CerIV, CerVA, and right CrusII) also presented a significant aging effect. Finally, to test for more fine grained effects of aging, the same sample was analyzed with VBM revealing additional clusters of GM that are significantly affected by age in chimpanzees (Figure 5) after FWE correction using threshold-free cluster enhancement (TFCE) (Smith and Nichols 2009). On top of the regions identified by region-wise morphometry, we found voxel-wise effects throughout anterior cingulate cortex (ACC), middle frontal gyrus (MFG) and in parts of the superior and inferior frontal gyrus (SFG, IFG), postcentral gyrus, superior and transverse temporal gyrus (STG, TTG), angular gyrus (AnG), superior occipital gyrus (sOG) and in inferior parts of the cerebellum.

**Figure 4.**
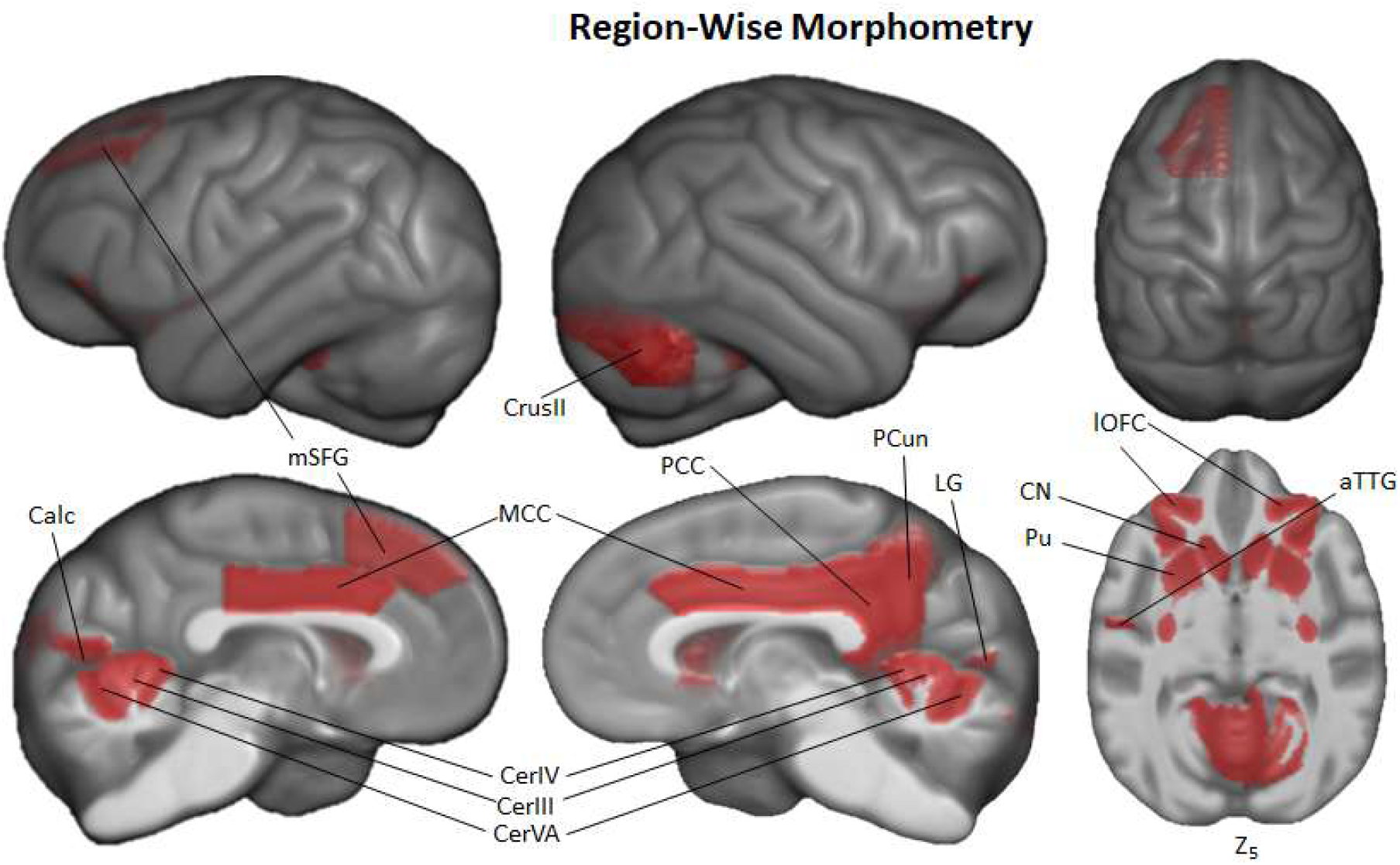
Region-wise morphometry in the Davi130 parcellation age regression where red regions represent Davi130 regions that remained significant at *p* ≤ 0.05 following FWE correction. aTTG – anterior transverse temporal gyrus, Calc – Calcarine sulcus, CrusII – cerebellum VIIA-CrusII, CerVA – cerebellum VA, CerIV – cerebellum IV, CerIII – cerebellum III, CN – caudate nucleus, lOFC – lateral orbitofrontal cortex, LG – lingual gyrus, MCC – medial cingulate cortex, mSFG – middle superior frontal gyrus, PCC – posterior cingulate cortex, PCun – precuneus, Pu – putamen.

**Figure 5.**
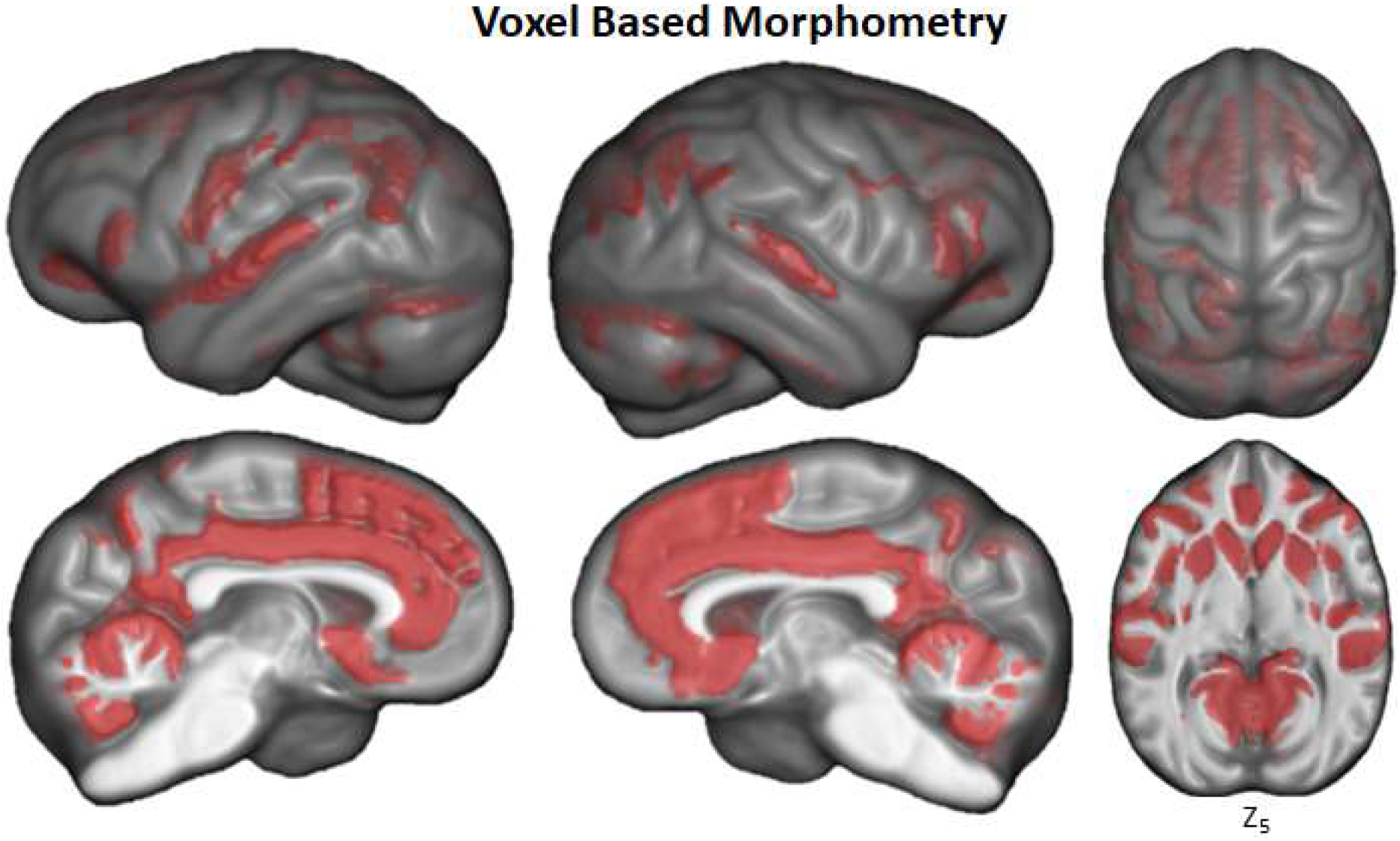
Voxel based morphometry of aging on GM volume using TFCE with FWE correction at *p* ≤ 0.05.

Hemispheric asymmetry of the chimpanzee brain was assessed for each cortical Davi130 region with a total of 29 macro-anatomical regions exhibiting significant cortical asymmetry after FWE correction (Figure 6, all region asymmetry found in Supplementary). Slightly more regions were found with greater GM volume on the right hemisphere (n=17) as compared to the left (n=12). In the left hemisphere, we found more GM laterally in the inferior postcentral gyrus (iPoCG), supramarginal gyrus (SMG) and superior occipital gyrus (sOG), insula and posterior TTG as well as medially in the orbitofrontal cortex (mOFG), basal forebrain nuclei (BF), hippocampus (HC), and anterior SFG. Rightward cortical asymmetry was located medially in the paracentral lobule (PCL), PCC, cuneus (Cun) and the region surrounding the calcarine sulcus, in the parahippocampal gyrus (PHC) and posterior fusiform gyrus (pFFG), as well as in the parietal operculum (POP). Laterally, rightward cortical asymmetry was found in the middle postcentral gyrus (mPoCG) and posterior superior temporal gyrus (pSTG). Within the basal ganglia, leftward GM asymmetry was observed in the putamen (Pu) while, rightward asymmetry is in the globus pallidus (GP) and thalamus (Th). In the anterior cerebellar lobe, there was leftward (CerIV) and rightward (CerII) GM asymmetry. The posterior cerebellum showed only rightward (CerVI, CerVB, and CerIX) asymmetry.

**Figure 6.**
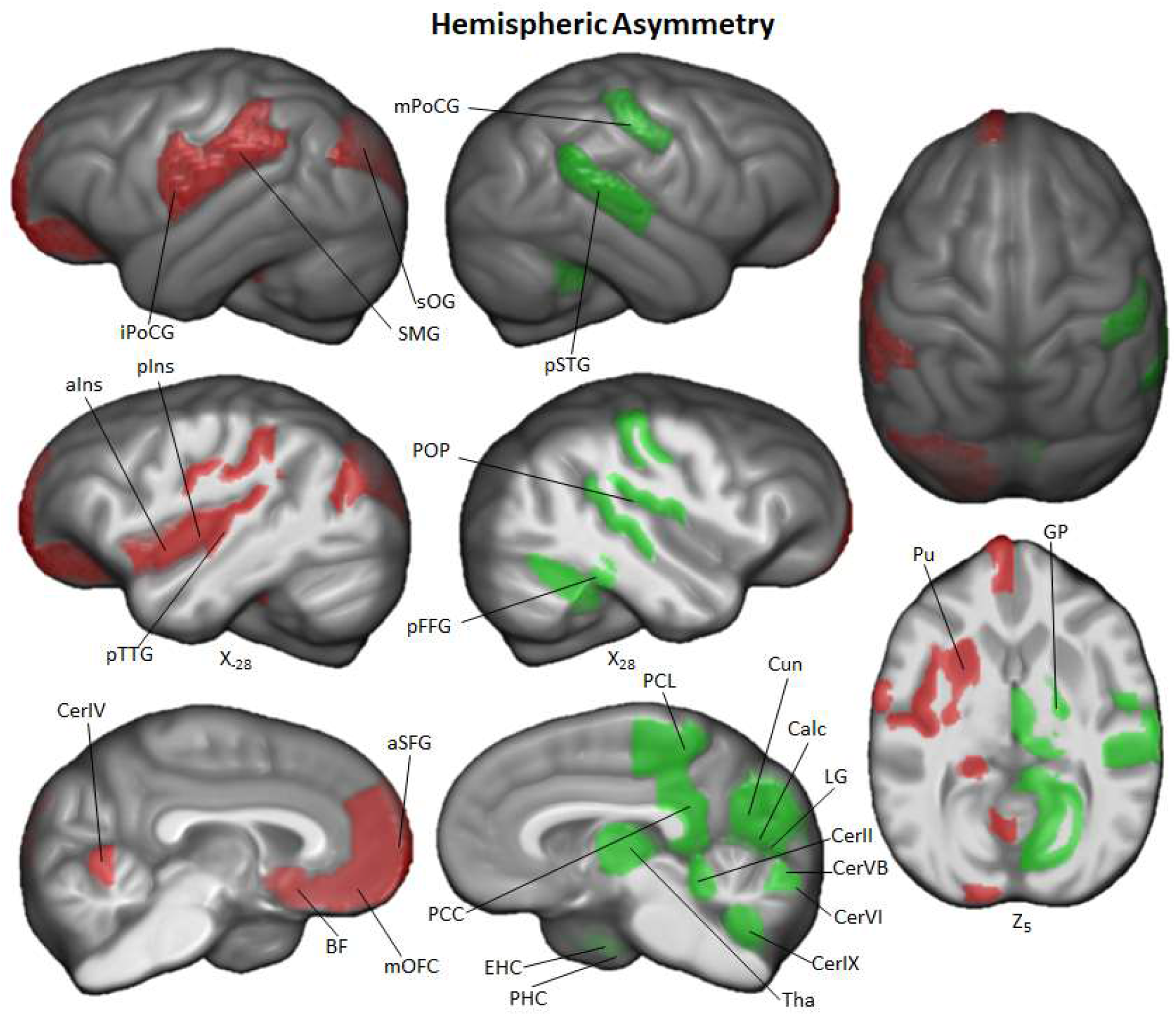
Hemispheric asymmetry of Davi130 regions within the chimpanzee sample. Significant leftward (red) and rightward (green) asymmetrical regions are those with a *p* ≤ 0.05 after FWE correction. aIns – anterior insula, aSFG – anterior superior frontal gyrus, BF – basal forebrain nuclei, Calc – calcarine sulcus, CerIX – cerebellum IX, CerVB – cerebellum V anterior B, CerVI – cerebellum VI, CerIV – cerebellum IV, CerII, cerebellum II, Cun – cuneus, EnC – entorhinal cortex, GP – globus pallidus, iPoCG – inferior postcentral gyrus, LG – lingual gyrus, mOFC – medial orbitofrontal cortex, mPoCG – middle postcentral gyrus, PCC – posterior cingulate cortex, PCL – paracentral lobule, pFFG – posterior fusiform gyrus, PHC – parahippocampal gyrus, pIns – posterior insula, POP – parietal operculum, pSTG – posterior superior temporal gyrus, pTTG – posterior transverse temporal gyrus, Pu – putamen, SMG – supramarginal gyrus, sOG – superior occipital gyrus, Th – thalamus. The hippocampus (HC) is significantly leftward lateralized but is not shown in this figure.

The moderate pattern of asymmetry we observed medially was a shift from more anterior regions showing leftward GM asymmetry (mOFC, BF, and aSFG) while more posterior regions presented rightward asymmetry (PCL, PCC, Cun, Calc, and LG). The lateral regions did not show a decipherable pattern.

## Discussion

As a common reference space for the analysis of chimpanzee brain data, we created the Juna.Chimp template, constructed from a large heterogeneous sample of T1-weighted MRI’s from the NCBR. The Juna.Chimp template includes a reference T1-template, along with probability maps of brain and head tissues accompanied by a Geodesic Shooting template for the publicly available SPM12/CAT12 preprocessing pipeline to efficiently segment and spatially normalize individual chimpanzee T1 images. The T1-template and TPM can be also used as the target for image registration with other popular software packages, such as FSL (https://fsl.fmrib.ox.ac.uk/fsl) or ANTs (http://stnava.github.io/ANTs/).

Additionally, we provide the manually segmented, macro-anatomical Davi130 whole-brain parcellation comprising 130 cortical, sub-cortical and cerebellar brain regions, which enables systematical extraction of volumes-of-interest from chimpanzee MRI data. The image processing pipeline and Davi130 parcellation were used to investigate ageing and interhemispheric asymmetry in the chimpanzee brain. Our analyses demonstrated strong age-related GM atrophy as well as marked hemispheric asymmetry over the whole cortex.

We found clear evidence of global and local GM decline in the aging chimpanzee brain even though previous research into age-related changes in chimpanzee brain organization has shown little to no effect (Herndon et al. 1999; Sherwood et al. 2011; Chen et al. 2013; Autrey et al. 2014). This can be attributed on the one hand to the larger number of MRI scans available via the NCBR including 30% of older subjects with 55 individuals over 30 and 12 over 45 years of age, which is crucial for modelling the effect of aging (Chen et al. 2013; Autrey et al. 2014). On the other hand, state-of-the-art image processing enabled the creation of the species specific Juna.Chimp templates, which largely improves tissue segmentation and registration accuracy (Ashburner and Friston 2000). Non-linear registration was also improved by the large heterogeneous sample utilized for the creation of the templates encompassing a representative amount of inter-individual variation. We used the well-established structural brain imaging toolbox CAT12 to build a reusable chimpanzee preprocessing pipeline catered towards analyzing local tissue-specific anatomical variations as measured with T1 weighted MRI. The Davi130-based region-wise and the voxel-wise morphometry analysis consistently showed localized GM decline in lOFC, the basal ganglia, MCC, PCC, PCun and superior cerebellum. The VBM approach additionally produced evidence for age effects in bilateral prefrontal cortex, ACC, superior temporal regions and throughout the cerebellum. These additional effects can be expected, as VBM is more sensitive to GM changes due to aging (Kennedy et al. 2009). The brain regions revealing GM decline in both approaches in particular medial and temporal cortical regions and the basal ganglia have also been shown to exhibit GM atrophy during healthy aging in humans (Good et al. 2001b; Kennedy et al. 2009; Crivello et al. 2014; Minkova et al. 2017).

Very recently, it has been shown that stress hormone levels increase with age in chimpanzees, a process previously thought to only occur in humans which can cause GM volume decline (Emery Thompson et al. 2020). This further strengthens the argument that age-related GM decline is also shared by humans closest relative, the chimpanzee. Furthermore, Edler et al. (2017) found Alzheimer’s disease-like accumulation of amyloid beta plaques and neurofibrillary tangles located predominantly in prefrontal and temporal cortices in a sample of elderly chimpanzees between 37 and 62 years of age. As the aggregation of these proteins is associated with localized neuronal loss and cortical atrophy in humans (La Joie et al. 2012; Llado et al. 2018), the age-related decline in GM volume shown here is well in line with the findings by Jagust (2016) associating GM atrophy with amyloid beta. These findings provide a biological mechanism for accelerated GM decrease in prefrontal, limbic, and temporal cortices in chimpanzees. In contrast, elderly rhesus monkeys show GM volume decline without the presence of neurofibrillary tangles (Alexander et al. 2008; Shamy et al. 2011). Taken together, regionally specific GM atrophy seems to be a common aspect of the primate brain aging pattern observed in macaque monkeys, chimpanzees and humans. Yet, to make a case for the existence of Alzheimer’s disease in chimpanzees, validated cognitive tests for Alzheimer’s-like cognitive decline in non-human primates are needed, to test for direct associations between cognitive decline with tau pathology and brain atrophy.

Hemispheric asymmetry was found in almost two-thirds of all regions of the Davi130 parcellation, reproducing several regional findings reported in previous studies using diverse image processing methods as well as uncovering numerous novel population-level asymmetries. Previous studies utilizing a region-wise approach based on hand-drawn or atlas derived regions also reported leftward asymmetry of volume of the planum temporale (PT) (Lyn et al. 2011; Gilissen and Hopkins 2013), and of cortical thickness in STG (Hopkins and Avants 2013), and the insula (Hopkins et al. 2017). Region-wise morphometry also demonstrated rightward asymmetry in thickness of the PCL and PHC (Hopkins et al. 2017). Previous VBM findings also revealed leftward asymmetry in the anterior SFG and SMG along with rightward lateralization of the posterior SFG, and middle part of the PoCG (Hopkins et al. 2008). In the current study, new regions of larger GM volume in the left hemisphere were found in frontal (mSFG, mOFC), limbic (HC), temporal (aTTG), and parietal (iPoCG) cortices as well as in the basal ganglia (BF, Pu) and cerebellum (CerIV). Novel rightward asymmetries could also be seen in temporal (pFFG), limbic (PCC, EnG), parietal (POP), and occipital (Cun, LG, Calc) cortices besides the basal ganglia (Th, GP) and the cerebellum (CerIX, CerVI, CerVB, CerII).

Significant leftward asymmetries in Davi130s’ region pTTG which contains the PT, is consistent with previous studies in GM volume of the PT, its surface area (Hopkins and Nir 2010), and cytoarchitecture (Zilles et al. 1996; Gannon et al. 1998; Spocter et al. 2010). At this cortical location, old world monkeys lack the morphological features of the PT, nevertheless several species have been shown to display asymmetry in Sylvian fissure length (Lyn et al. 2011; Marie et al. 2018).

Interestingly, the parietal operculum showed rightward asymmetry, while Gilissen and Hopkins (2013) showed that the left parietal operculum was significantly longer in chimpanzees, compared to the right. The left lateral sulcus in the Juna.Chimp template appears to proceed further posteriorly and superiorly compared to the right, confirming this finding, even though we found greater GM volume in the right POP as compared to the left.

Population-level asymmetries in the pIFG in chimpanzees were documented almost two decades ago by Cantalupo and Hopkins (2001), who reported a leftward asymmetry in pIFG volume in a small sample of great apes. In subsequent studies this result could not be replicated when considering GM volume (Hopkins et al. 2008; Keller et al. 2009) or cytoarchitecture (Schenker et al. 2010). We also failed to find a leftward asymmetry in GM volume for the pIFG, in contrary to asymmetries found in humans (Amunts et al. 1999; Uylings et al. 2006; Keller et al. 2009). The prominent leftward PT asymmetry in chimpanzees is also a well-documented population-level asymmetry in humans (Good et al. 2001a; Watkins 2001). The overall regional distribution of asymmetry in chimpanzees is partially similar to that found in human cortical organization (Good et al. 2001a; Luders et al. 2006; Zhou et al. 2013; Koelkebeck et al. 2014; Plessen et al. 2014; Chiarello et al. 2016; Maingault et al. 2016; Kong et al. 2018).

Gross hemispheric asymmetry in humans follows a general structure of frontal rightward and occipital leftward asymmetry known as the ‘Yakovlevian torque’. This refers to the bending of the anterior right hemisphere over the midline into the left and the posterior left hemisphere bending over to the right and is represented as differences in widths of frontal and occipital lobes (Toga and Thompson 2003). This organizational trait was apparent in the Juna.Chimp templates with a slight frontal and occipital bending, which was manually adapted when labelling the medial Davi130 regions. The higher leftward GM density in frontal regions and rightward asymmetry in occipital regions may also follow this trend. Of note, this organizational pattern of asymmetry was less apparent in lateral cortical structures, however medially, this anterior-posterior asymmetry is evident in chimpanzees, challenging recent findings from Li and colleagues (2018) who rejected the ‘Yakovlevian torque’ in chimpanzees. Specifically, leftward asymmetry of the frontal regions aSFG, BF, and mOFC and the rightward asymmetry of posterior regions PCL, PCC, Cun, Calc, and LG aligns with the pattern of asymmetry reported in human cortical GM volume, thickness and surface area (Luders et al. 2006; Zhou et al. 2013; Plessen et al. 2014; Chiarello et al. 2016; Kong et al. 2018).

The NCBR offers the largest and richest openly available dataset of chimpanzee brain MRI scans acquired over a decade with 1.5T and 3T MRI at two locations, capturing valuable inter-individual variation in one large heterogeneous sample. To account for the scanner effect on GM estimation, field strength was modelled as a covariate of no interest for analyzing the age effect on GM volume. Rearing has been shown to affect GM structural covariance networks and cortical organization (Bogart et al. 2014; Bard and Hopkins 2018) while handedness has been shown to correlate with asymmetry in the motor cortex (Hopkins and Cantalupo 2004) and the volume of IFG (Taglialatela et al. 2006) as well as with gyrification asymmetry (Hopkins et al. 2007). Therefore, these covariates may also modulate hemispheric asymmetry and/or age-related GM volume decline here, but as these data were not available for all subjects, we did not consider these effects in order to include as many subjects as possible to model effects of age. The focus of this study was the analysis of GM volume, even though the CAT12 image processing pipeline includes surface projection and analysis. Consequently, the next step will be the application of CAT12 to analyze cortical surface area, curvature, gyrification, and thickness of the chimpanzee brain, to include behavioral data and the quantitative comparison to humans and other species, as cortical surface projection permits a direct inter-species comparison due to cross species registration.

## Conclusion

In conclusion, we present the new chimpanzee reference template Juna.Chimp, TPM’s, the Davi130 whole brain parcellation, and the CAT12 preprocessing pipeline which is ready-to-use by the wider neuroimaging community. Investigations of age-related GM changes in chimpanzees using both region-wise and voxel-based morphometry, showed a substantial age effect, providing further evidence for human like physiological aging processes in chimpanzees. Examining population based cortical asymmetry in chimpanzees yielded further evidence for the well-documented lateralization of PT. Additionally, an anterior-posterior left-right pattern of asymmetry as observed in humans was found predominantly in medial regions of the chimpanzee cortex.

## Materials and Methods

### Subject Information and Image Collection Procedure

This study analyzed structural T1-weighted MRI scans of 223 chimpanzees (137 females; 9 - 54 y/o, mean age 26.9 ± 10.2 years) from the NCBR (www.chimpanzeebrain.org). The chimpanzees were housed at two locations including, the *National Center for Chimpanzee Care of The University of Texas MD Anderson Cancer Center* (UTMDACC) and the *Yerkes National Primate Research Center* (YNPRC) of Emory University. The standard MR imaging procedures for chimpanzees at the YNPRC and UTMDACC are designed to minimize stress for the subjects. For an in-depth explanation of the imaging procedure please refer to Autrey et al. (2014). Seventy-six chimpanzees were scanned with a Siemens Trio 3 Tesla scanner (Siemens Medical Solutions USA, Inc., Malvern, Pennsylvania, USA). Most T1-weighted images were collected using a threedimensional gradient echo sequence with 0.6 × 0.6 × 0.6 resolution (pulse repetition = 2300 ms, echo time = 4.4 ms, number of signals averaged = 3). The remaining 147 chimpanzees were scanned using a 1.5T GE echo-speed Horizon LX MR scanner (GE Medical Systems, Milwaukee, WI), predominantly applying gradient echo sequence with 0.7 × 0.7 × 1.2 resolution (pulse repetition = 19.0 ms, echo time = 8.5 ms, number of signals averaged = 8).

### Creation of Chimpanzee Templates

An iterative process as by Franke et al. (2017) was employed to create the Juna.Chimp template, with T1 average, Shooting registration template (Ashburner and Friston 2011), as well as the TPM (Figure 7). Initially, a first-generation template was produced using the “greater_ape” template delivered by CAT (Dahnke and Gaser 2017; Gaser et al. 2020) that utilizes data provided in Rilling and Insel (1999). The final segmentation takes the bias-corrected, intensity-normalized, and skull-stripped image together with the initial SPM-segmentation to conduct an Adaptive Maximum A Posterior (AMAP) estimation (Rajapakse et al. 1997) with a partial volume model for sub-voxel accuracy (Tohka et al. 2004). The affine normalized tissue segments of GM, WM, and CSF were used to create a new Shooting template that consists of four major non-linear normalization steps allowing to normalize new scans. To create a chimpanzee-specific TPM, we average the different Shooting template steps to benefit from the high spatial resolution of the final Shooting steps but also include the general affine aspects to avoid overoptimization. Besides the brain tissues the TPM also included two head tissues (bones and muscles) and a background class for standard SPM12 (Ashburner and Friston 2005) and CAT12 preprocessing. The internal CAT atlas was written for each subject and mapped to the new chimpanzee template using the information from the Shooting registration. The CAT atlas maps were averaged by a median filter and finally manually corrected. This initial template was then used in the second iteration of CAT segmentation to establish the final chimpanzee-specific Juna.Chimp template, which was imported into the standard CAT12 preprocessing pipeline to create the final data used for the aging and asymmetry analyses.

**Figure 7.**
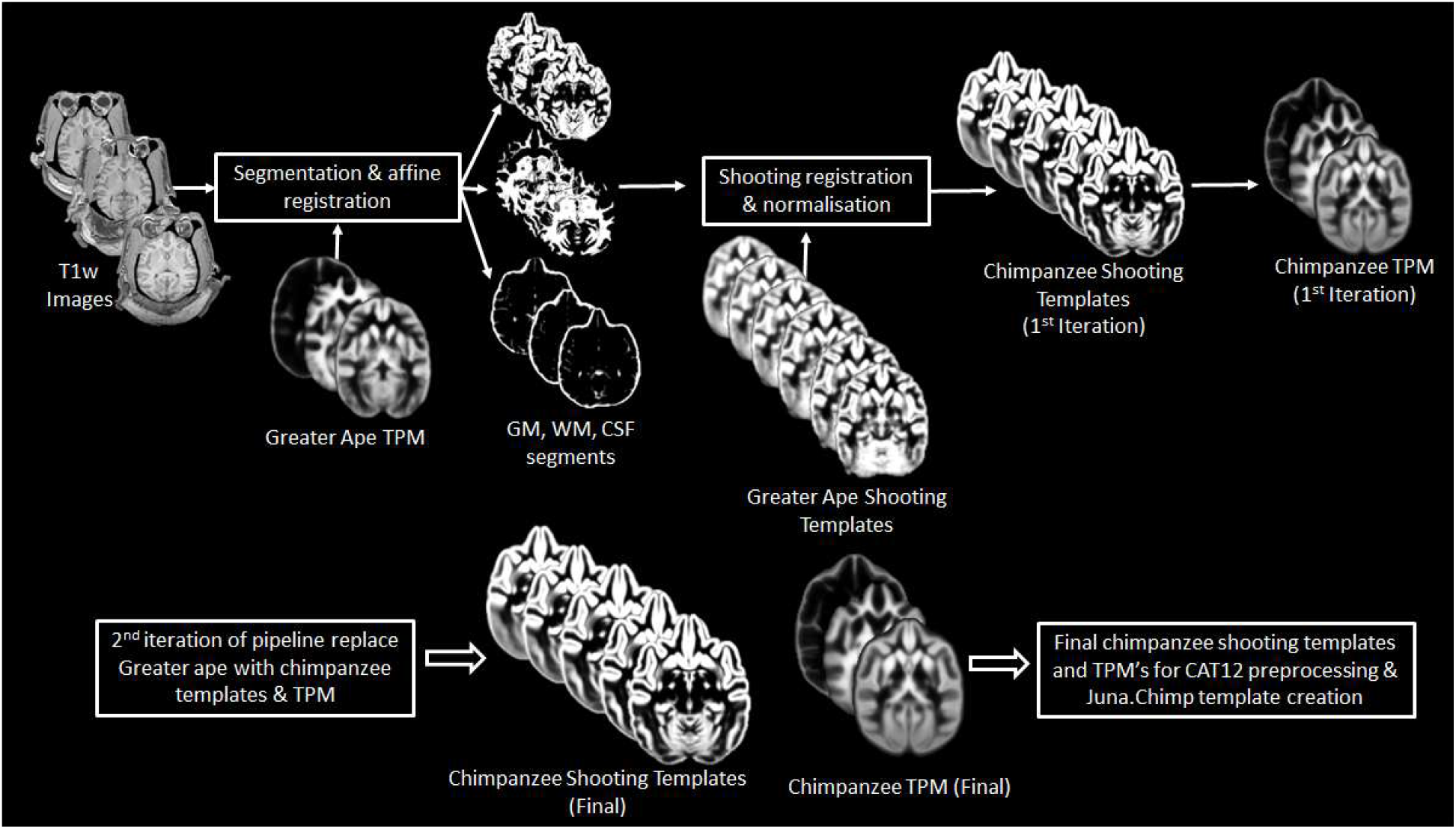
Workflow for creation of chimpanzee specific Shooting template and TPM, which can then be used in CAT12 structural preprocessing pipeline to create the Juna.Chimp template.

The resulting chimpanzee-Shooting template, TPM and CAT atlas establishes the robust and reliable base to segment and spatially normalize the T1-weighted images utilizing CAT12’s processing pipeline (Dahnke and Gaser 2017; Gaser et al. 2020).

### Davi130 parcellation

The average T1 and final Shooting template were used for a manual delineation of macro-anatomical GM structures. Identification and annotation of major brain regions were performed manually using the program, 3D Slicer 4.10.1 (https://www.slicer.org). The labeling enables automated, region-based analysis of the entire chimpanzee brain and allows for robust statistical analysis. Nomenclature and location of regions were ascertained by consulting both chimpanzee and human brain atlases (Bailey P, Bonin GV 1950; Mai et al. 2015). The labelling was completed by two authors (S.V. & R.D.) and reviewed by two experts of chimpanzee brain anatomy (C.C.S. & W.D.H.).

A total of 65 GM structures within the cerebrum and cerebellum of the left hemisphere were annotated and then flipped to the right hemisphere. The flipped annotations were then manually adapted to the morphology of the right hemisphere to have complete coverage of the chimpanzee brain with 130 labels. The slight bending of the anterior part of the right hemisphere and the posterior part of the left hemisphere over the midline observed in our template and annotated within our Davi130 labelling, does not align with previous findings claiming that this morphological trait is specific to the human brain (Li et al. 2018; Xiang et al. 2019).

The location of macroscopic brain regions was determined based on major gyri of the cerebral cortex, as well as distinct anatomical landmarks of the cerebellar cortex, and basal ganglia. Of note, the border between two gyri was arbitrarily set as the mid-point of the connecting sulcus. Large gyri were further subdivided into two or three parts based on their size and structural features, such as sulcal fundi and gyral peaks, to enable greater spatial resolution and better inter-regional comparison. Naming of subdivisions was based entirely on spatial location, e.g., anterior, middle, posterior, and do not claim to correspond to functional parcellations.

### Quality Control

Rigorous QC was employed on all images using two iterative steps. The first step utilized the built-in CAT12 quality assurance and ‘check sample homogeneity’ function. The modulated GM maps were initially tested for sample in-homogeneity for each scanner strength separately (1.5T and 3T). The images that passed the first QC step went through a final round of sample inhomogeneity testing as a whole sample to finally arrive at our study sample, which included 178 chimpanzees (120 females, 11 – 54 years old, mean = 26.7 ± 9.8). A more in-depth explanation of the QC procedure can be found in Supplementary 1.2.

### Age-Related Changes in Total Gray Matter

A linear regression model was used to determine the effect of aging on total GM volume. Firstly, total GM volume for each subject was converted into a percentage of total intracranial volume (TIV) to account for the variation in head size. This was then entered into a linear regression model as the dependent variable with age and sex as the independents. Sex-specific models were conducted with males and females separately using age as the only dependent variable. The slope of each regression line was determined using R^2^ and a p-value of *p* ≤ 0.05 was used to determine the significant effect of age and sex on total GM volume.

### Age-Related Changes in Gray Matter Using Davi130 Parcellation

The Davi130 parcellation was applied to the modulated GM maps to conduct region-wise morphometry analysis. First, the Davi130 regions were masked with a 0.1 GM mask to remove all non-GM portions of the regions. Subsequently, the average GM intensity of each region for all QC-passed chimpanzees was calculated. A multiple regression model was conducted for the labels from both hemispheres, whereby, the dependent variable was GM volume and the predictor variables were age, sex, TIV, and scanner. Significant age-related GM decline was established for a Davi130 label with a *p* ≤ 0.05, after correcting for multiple comparisons using FWE (Holm 1979).

### Voxel-Based Morphometry

VBM analysis was conducted using CAT12 to determine the effect of aging on local GM volume. The modulated and spatially normalized GM segments from each subject were spatially smoothed with a 4 mm FWHM (full width half maximum) kernel prior to analyses. To restrict the overall volume of interest, an implicit 0.4 GM mask was employed. As MRI field strength is known to influence image quality, and consequently, tissue classification, we included scanner strength in our VBM model as a covariate. The dependent variable in the model was age, with covariates of TIV, sex, and scanner. The VBM model was corrected for multiple comparisons using TFCE with 5000 permutations (Smith and Nichols 2009). Significant clusters were determined at *p* ≤ 0.05, after correcting for multiple comparisons using FWE.

### Hemispheric Asymmetry

As for the age regression analysis, all Davi130 parcels were masked with a 0.1 GM mask to remove non-GM portions within regions. Cortical hemispheric asymmetry of Davi130 labels was determined using the formula *Asym* = *(L − R) / (L + R) * 0.5* (Kurth et al. 2015; Hopkins et al. 2017), whereby L and R represent the average GM volume for each region in the left and right hemisphere, respectively. Therefore, the bi-hemispheric Davi130 regions were converted into single *Asym* labels (n=65) with positive *Asym* values indicating a leftward asymmetry, and negative values, a rightward bias. One-sample *t*-tests were conducted for each region under the null hypothesis of *Asym* = *0*, and significant leftward or rightward asymmetry was determined with a *p* ≤ 0.05, after correcting for multiple comparisons using FWE (Holm 1979).

### Exemplar Pipeline Workflow

To illustrate the structural processing pipeline, we have created exemplar MATLAB SPM batch scripts that utilizes the Juna.Chimp templates in CAT12’s preprocessing workflow to conduct segmentation, spatial registration, and finally some basic age analysis on an openly available direct-to-download chimpanzee sample (http://www.chimpanzeebrain.org/mri-datasets-for-direct-download). These scripts require the appropriate templates which can be downloaded from the Juna.Chimp web viewer (SPM/CAT_templates.zip) and then place the templates_animals/ folder into the latest version CAT12 Toolbox directory (CAT12.7 r1609). The processing parameters are similar to those conducted in this study, although different DICOM conversions and denoising were conducted. Further information regarding each parameter can be viewed when opening the script in the SPM batch as well as the provided comments and README file. The code for the workflow in addition to the code used to conduct the aging effect and asymmetry analyses can be found here https://github.com/viko18/JunaChimp.

## Acknowledgements

We would like to thank Jona Fischer for the creation of the interactive Juna.Chimp web viewer adapted from nehuba (github). This study was supported by the European Union’s Horizon 2020 Research and Innovation Program under Grant Agreement No. 785907 (HBP SGA2).

## Supplementary

### Supplementary 1.1 DICOM conversion and De-noising

The structural T1-weighted images were provided by the NCBR in their original DICOM format and converted into Nifti using MRIcron (Rorder and Brett 2000). If multiple scans were available, the average was computed. Following DICOM conversion each image was cleaned of noise (Manjon et al. 2010) and signal inhomogeneity and resliced to 0.6 mm isotropic resolution. Finally, the anterior commissure was manually set as the center (0,0,0) of all Nifti’s to aid in affine preprocessing.

**Supplementary Figure 1.**
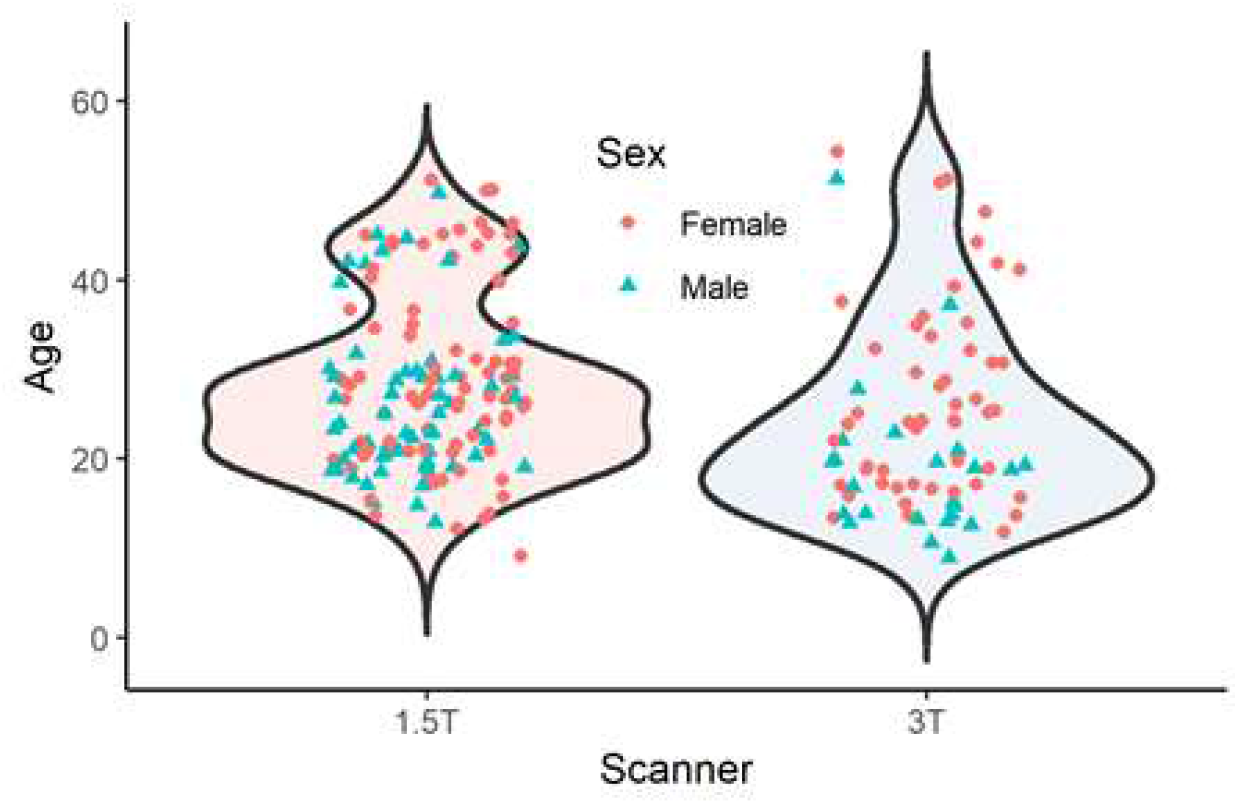
Age and sex distribution of complete 223 chimpanzees separated by scanner strength.

### Supplementary 1.2 Chimpanzee QC

CAT12 provides quality measures pertaining to noise, bias inhomogeneities, resolution and an overall compounded score of the original input image. Using these ratings, poor images were flagged for visual inspection when they were 2 standard deviations (std) away from the sample mean of each rating. The preprocessed modulated GM maps were then tested for sample inhomogeneity separately for each scanner (3T & 1.5T) and those that have a mean correlation below 2 std were flagged for visual inspection. Once the original image was flagged, affine GM, and modulated GM maps were inspected for poor quality, tissue misclassification, artefacts, irregular deformations, and very high intensities. For the second iteration, the passed modulated GM maps were tested again for mean correlation as a complete sample, flagging the images below 3 std for visual inspection. Looking for the same features as in the initial QC iteration. Following the two iterations of QC a total of 178 of 223 chimpanzee MRI’s (120 females, 11 – 54 y/o, mean = 26.7 ± 9.8) qualified for statistical analysis.

### Supplementary 1.3 CAT12 Preprocessing Segmentation

Structural image segmentation in CAT12 builds on the TPM-based approach employed by SPM12, whereby, the gray/white image intensity is aided with a priori tissue probabilities in initial segmentation and affine registration as it is in common template space. Another advantage of a TPM is that one has a template for initial affine registration, which then enables the segment maps to be non-linearly registered and spatially normalized to corresponding segment maps of the chimpanzee shooting templates.

Lowering the possibility for registration errors improves the quality of the final normalized image. Improving upon SPM’s segmentation (Ashburner and Friston 2005), CAT12 employs Local Adaptive Segmentation (LAS) (Dahnke et al. 2012), Adaptive Maximum A Posterior segmentation(AMAP) (Dahnke and Gaser 2017; Gaser et al. 2020), and Partial Volume Estimation (PVE) (Tohka et al. 2004). LAS creates local intensity transformations for all tissue types to limit GM misclassification due to varying GM intensity in regions such as the occipital, basal ganglia, and motor cortex as a result of anatomical properties (e.g. high myelination and iron content). AMAP segmentation takes the initially segmented, aligned, and skull stripped image created utilizing the TPM and disregards the a priori information of the TPM, to conduct an adaptive AMAP estimation where local variations are modelled by slowly varying spatial functions (Rajapakse et al. 1997). Along with the classical three tissue types for segmentation (GM, WM, & CSF) based on the AMAP estimation, an additional two PVE classes (GM-WM & GM-CSF) are created resulting in an estimate of the fraction of each tissue type contained in each voxel. These features outlined above of our pipeline allow for more accurate tissue segmentation and therefore a better representation of macroanatomical GM levels for analysis.

**Supplementary Table 1.** Davi130 Labels

| Label Number | Davi130 Label | Brain Region | Hemisphere | Acronyms |
| --- | --- | --- | --- | --- |
| 1 | Anterior Superior Frontal Gyrus | Frontal | left | L.aSFG |
| 2 | Anterior Superior Frontal Gyrus | Frontal | right | R.aSFG |
| 3 | Middle Superior Frontal Gyrus | Frontal | left | L.mSFG |
| 4 | Middle Superior Frontal Gyrus | Frontal | right | R.mSFG |
| 5 | Posterior Superior Frontal Gyrus | Frontal | left | L.Hth |
| 6 | Posterior Superior Frontal Gyrus | Frontal | right | R.pSFG |
| 7 | Anterior Middle Frontal Gyrus | Frontal | left | L.aMFG |
| 8 | Anterior Middle Frontal Gyrus | Frontal | right | R.aMFG |
| 9 | Posterior Middle Frontal Gyrus | Frontal | left | L.pMFG |
| 10 | Posterior Middle Frontal Gyrus | Frontal | right | R.pMFG |
| 11 | Anterior Inferior Frontal Gyrus | Frontal | left | L.aIFG |
| 12 | Anterior Inferior Frontal Gyrus | Frontal | right | R.aIFG |
| 13 | Middle Inferior Frontal Gyrus | Frontal | left | L.mIFG |
| 14 | Middle Inferior Frontal Gyrus | Frontal | right | R.mIFG |
| 15 | Posterior Inferior Frontal Gyrus | Frontal | left | L.pIFG |
| 16 | Posterior Inferior Frontal Gyrus | Frontal | right | R.pIFG |
| 17 | Medial Orbitofrontal Cortex | Frontal | left | L.mOFC |
| 18 | Medial Orbitofrontal Cortex | Frontal | right | R.mOFC |
| 19 | Lateral Orbitofrontal Cortex | Frontal | left | L.lOFC |
| 20 | Lateral Orbitofrontal Cortex | Frontal | right | R.lOFC |
| 21 | Anterior Cingulate Cortex | Limbic | left | L.ACC |
| 22 | Anterior Cingulate Cortex | Limbic | right | R.ACC |
| 23 | Middle Cingulate Cortex | Limbic | left | L.MCC |
| 24 | Middle Cingulate Cortex | Limbic | right | R.MCC |
| 25 | Posterior Cingulate Cortex | Limbic | left | L.PCC |
| 26 | Posterior Cingulate Cortex | Limbic | right | R.PCC |
| 27 | Superior Precentral Gyrus | Frontal | left | L.sPrCG |
| 28 | Superior Precentral Gyrus | Frontal | right | R.sPrCG |
| 29 | Middle Precentral Gyrus | Frontal | left | L.mPrCG |
| 30 | Middle Precentral Gyrus | Frontal | right | R.mPrCG |
| 31 | Inferior Precentral Gyrus | Frontal | left | L.iPrCG |
| 32 | Inferior Precentral Gyrus | Frontal | right | R.iPrCG |
| 33 | Paracentral Lobule | Parietal | left | L.PCL |
| 34 | Paracentral Lobule | Parietal | right | R.PCL |
| 35 | Frontal Operculum | Frontal | left | L.FOP |
| 36 | Frontal Operculum | Frontal | right | R.FOP |
| 37 | Parietal Operculum | Parietal | left | L.POP |
| 38 | Parietal Operculum | Parietal | right | R.POP |
| 39 | Anterior Insula | Temporal | left | L.aIns |
| 40 | Anterior Insula | Temporal | right | R.aIns |
| 41 | Posterior Insula | Temporal | left | L.pIns |
| 42 | Posterior Insula | Temporal | right | R.pIns |
| 43 | Anterior Transverse Temporal Gyrus | Temporal | left | L.aTTG |
| 44 | Anterior Transverse Temporal Gyrus | Temporal | right | R.aTTG |
| 45 | Posterior Transverse Temporal Gyrus | Temporal | left | L.pTTG |
| 46 | Posterior Transverse Temporal Gyrus | Temporal | right | R.pTTG |
| 47 | Anterior Superior Temporal Gyrus | Temporal | left | L.aSTG |
| 48 | Anterior Superior Temporal Gyrus | Temporal | right | R.aSTG |
| 49 | Posterior Superior Temporal Gyrus | Temporal | left | L.pSTG |
| 50 | Posterior Superior Temporal Gyrus | Temporal | right | R.pSTG |
| 51 | Anterior Middle Temporal Gyrus | Temporal | left | L.aMTG |
| 52 | Anterior Middle Temporal Gyrus | Temporal | right | R.aMTG |
| 53 | Posterior Middle Temporal Gyrus | Temporal | left | L.pMTG |
| 54 | Posterior Middle Temporal Gyrus | Temporal | right | R.pMTG |
| 55 | Anterior Inferior Temporal Gyrus | Temporal | left | L.aITG |
| 56 | Anterior Inferior Temporal Gyrus | Temporal | right | R.aITG |
| 57 | Posterior Inferior Temporal Gyrus | Temporal | left | L.pITG |
| 58 | Posterior Inferior Temporal Gyrus | Temporal | right | R.pITG |
| 59 | Entorhinal Cortex | Limbic | left | L.EHC |
| 60 | Entorhinal Cortex | Limbic | right | R.EHC |
| 61 | Anterior Fusiform Gyrus | Temporal | left | L.aFFG |
| 62 | Anterior Fusiform Gyrus | Temporal | right | R.aFFG |
| 63 | Posterior Fusiform Gyrus | Temporal | left | L.pFFG |
| 64 | Posterior Fusiform Gyrus | Temporal | right | R.pFFG |
| 65 | Parahippocampal Gyrus | Limbic | left | L.PHC |
| 66 | Parahippocampal Gyrus | Limbic | right | R.PHC |
| 67 | Amygdala | Limbic | left | L.Amy |
| 68 | Amygdala | Limbic | right | R.Amy |
| 69 | Hippocampus | Limbic | left | L.HC |
| 70 | Hippocampus | Limbic | right | R.HC |
| 71 | Superior Postcentral Gyrus | Parietal | left | L.sPoCG |
| 72 | Superior Postcentral Gyrus | Parietal | right | R.sPoCG |
| 73 | Middle Postcentral Gyrus | Parietal | left | L.mPoCG |
| 74 | Middle Postcentral Gyrus | Parietal | right | R.mPoCG |
| 75 | Inferior Postcentral Gyrus | Parietal | left | L.iPoCG |
| 76 | Inferior Postcentral Gyrus | Parietal | right | R.iPoCG |
| 77 | Superior Parietal Lobule | Parietal | left | L.SPL |
| 78 | Superior Parietal Lobule | Parietal | right | R.SPL |
| 79 | Supramarginal Gyrus | Parietal | left | L.SMG |
| 80 | Supramarginal Gyrus | Parietal | right | R.SMG |
| 81 | Angular Gyrus | Parietal | left | L.AnG |
| 82 | Angular Gyrus | Parietal | right | R.AnG |
| 83 | Precuneus | Parietal | left | L.PCun |
| 84 | Precuneus | Parietal | right | R.PCun |
| 85 | Cuneus | Occipital | left | L.Cun |
| 86 | Cuneus | Occipital | right | R.Cun |
| 87 | Lingual Gyrus | Occipital | left | L.LG |
| 88 | Lingual Gyrus | Occipital | right | R.LG |
| 89 | Calcarine Sulcus | Occipital | left | L.Calc |
| 90 | Calcarine Sulcus | Occipital | right | R.Calc |
| 91 | Superior Occipital Gyrus | Occipital | left | L.sOG |
| 92 | Superior Occipital Gyrus | Occipital | right | R.sOG |
| 93 | Middle Occipital Gyrus | Occipital | left | L.mOG |
| 94 | Middle Occipital Gyrus | Occipital | right | R.mOG |
| 95 | Inferior Occipital Gyrus | Occipital | left | L.iOG |
| 96 | Inferior Occipital Gyrus | Occipital | right | R.iOG |
| 97 | Caudate Nucleus | Basal Ganglia | left | L.CN |
| 98 | Caudate Nucleus | Basal Ganglia | right | R.CN |
| 99 | Nucleus Accumbens | Basal Ganglia | left | L.NA |
| 100 | Nucleus Accumbens | Basal Ganglia | right | R.NA |
| 101 | Basal Forebrain Nuclei | Basal Ganglia | left | L.BF |
| 102 | Basal Forebrain Nuclei | Basal Ganglia | right | R.BF |
| 103 | Putamen | Basal Ganglia | left | L.Pu |
| 104 | Putamen | Basal Ganglia | right | R.Pu |
| 105 | Globus pallidus | Basal Ganglia | left | L.GP |
| 106 | Globus pallidus | Basal Ganglia | right | R.GP |
| 107 | Thalamus | Basal Ganglia | left | L.Th |
| 108 | Thalamus | Basal Ganglia | right | R.Th |
| 109 | Hypothalamus | Basal Ganglia | left | L.HTh |
| 110 | Hypothalamus | Basal Ganglia | right | R.HTh |
| 111 | Cerebellum IX-Tonsil | Cerebellum | left | L.CerIX |
| 112 | Cerebellum IX-Tonsil | Cerebellum | right | R.CerIX |
| 113 | Cerebellum VIIIAB-Inferior Posterior - PML | Cerebellum | left | L.CerVIII |
| 114 | Cerebellum VIIIAB-Inferior Posterior - PML | Cerebellum | right | R.CerVIII |
| 115 | Cerebellum VIIA - Superior Posterior - Crus I of Ansiform Lobule | Cerebellum | left | L.CrusI |
| 116 | Cerebellum VIIA - Superior Posterior – Crus I of Ansiform Lobule | Cerebellum | right | R.CrusI |
| 117 | Cerebellum VIIA - Superior Posterior – Crus II of Ansiform Lobule with Paramedian 1 | Cerebellum | left | L.CrusII |
| 118 | Cerebellum VIIA-Superior Posterior - Crus II of Ansiform Lobule with Paramedian 1 | Cerebellum | right | R.CrusII |
| 119 | Cerebellum VI-Superior Posterior | Cerebellum | left | L.CerVI |
| 120 | Cerebellum VI-Superior Posterior | Cerebellum | right | R.CerVI |
| 121 | Cerebellum V-Anterior B | Cerebellum | left | L.CerVB |
| 122 | Cerebellum V-Anterior B | Cerebellum | right | R.CerVB |
| 123 | Cerebellum V-Anterior A | Cerebellum | left | L.CerVA |
| 124 | Cerebellum V-Anterior A | Cerebellum | right | R.CerVA |
| 125 | Cerebellum IV-Anterior Quadrangulate | Cerebellum | left | L.CerIV |
| 126 | Cerebellum IV-Anterior Quadrangulate | Cerebellum | right | R.CerIV |
| 127 | Cerebellum III-Anterior Quadrangulate | Cerebellum | left | L.CerIII |
| 128 | Cerebellum III-Anterior Quadrangulate | Cerebellum | right | R.CerIII |
| 129 | Cerebellum II-Anterior Quadrangulate | Cerebellum | left | L.CerII |
| 130 | Cerebellum II-Anterior Quadrangulate | Cerebellum | right | R.CerII |

**Supplementary Table 2.** DaVi Labels Age Effect on Gray Matter Volume

| ***Davi130 Label*** | ***T-statistic*** | ***p-value*** |
| --- | --- | --- |
| **Frontal Cortex** |  |  |
| Anterior Superior Frontal Gyrus (L.aSFG) | -1.57 | 0.1197 |
| Anterior Superior Frontal Gyrus (R.aSFG) | -1.16 | 0.2482 |
| Middle Superior Frontal Gyrus (L.mSFG)* | -3.59 | 0.0004 |
| Middle Superior Frontal Gyrus (R.mSFG) | -3.55 | 0.0005 |
| Posterior Superior Frontal Gyrus (L.pSFG) | -3.18 | 0.0017 |
| Posterior Superior Frontal Gyrus (R.pSFG) | -2.35 | 0.0199 |
| Anterior Middle Frontal Gyrus (L.aMFG) | -1.08 | 0.2814 |
| Anterior Middle Frontal Gyrus (R.aMFG) | -2.11 | 0.0365 |
| Posterior Middle Frontal Gyrus (L.pMFG) | -1.88 | 0.0614 |
| Posterior Middle Frontal Gyrus (R.pMFG) | -1.79 | 0.0757 |
| Anterior Inferior Frontal Gyrus (L.aIFG) | -1.27 | 0.2048 |
| Anterior Inferior Frontal Gyrus (R.aIFG) | -1.52 | 0.1300 |
| Middle Inferior Frontal Gyrus (L.mIFG) | -1.79 | 0.0747 |
| Middle Inferior Frontal Gyrus (R.mIFG) | -2.65 | 0.0089 |
| Posterior Inferior Frontal Gyrus (L.pIFG) | -2.83 | 0.0052 |
| Posterior Inferior Frontal Gyrus (R.pIFG) | -2.00 | 0.0468 |
| Medial Orbitofrontal Cortex (L.mOFC) | -1.82 | 0.0711 |
| Medial Orbitofrontal Cortex (R.mOFC) | -1.83 | 0.0697 |
| Lateral Orbitofrontal Cortex (L.IOFC)* | -4.31 | 3.0x10<sup>-5</sup> |
| Lateral Orbitofrontal Cortex (R.IOFC)* | -3.79 | 0.0002 |
| Superior Precentral Gyrus (L.sPrCG) | -1.96 | 0.0511 |
| Superior Precentral Gyrus (R.sPrCG) | -0.47 | 0.6380 |
| Middle Precentral Gyrus (L.mPrCG) | -1.96 | 0.0519 |
| Middle Precentral Gyrus (R.mPrCG) | -1.34 | 0.1836 |
| Inferior Precentral Gyrus (L.iPrCG) | -1.78 | 0.0766 |
| Inferior Precentral Gyrus (R.iPrCG) | -1.20 | 0.2312 |
| Frontal Operculum (L.FOP) | -2.74 | 0.0067 |
| Frontal Operculum (R.FOP) | -1.55 | 0.1227 |
| **Limbic Cortex** |  |  |
| Anterior Cingulate Gyrus (L.ACC) | -2.66 | 0.0085 |
| Anterior Cingulate Gyrus (R.ACC) | -3.08 | 0.0024 |
| Middle Cingulate Gyrus (L.MCC)* | -4.02 | 8.78x10<sup>-5</sup> |
| Middle Cingulate Gyrus (R.MCC)* | -4.29 | 2.99x10<sup>-5</sup> |
| Posterior Cingulate Gyrus (L.PCC) | -3.37 | 0.0009 |
| Posterior Cingulate Gyrus (R.PCC)* | -3.81 | 0.0002 |
| Entorhinal Cortex (L.EG) | -0.43 | 0.6685 |
| Entorhinal Cortex (R.EG) | -1.22 | 0.2229 |
| Parahippocampal Gyrus (L.PHC) | -2.70 | 0.0077 |
| Parahippocampal Gyrus (R.PHC) | -3.15 | 0.0019 |
| Amygdala (L.Amy) | -3.26 | 0.0014 |
| Amygdala (R.Amy) | -2.73 | 0.0070 |
| Hippocampus (L.HC) | -0.72 | 0.4735 |
| Hippocampus (R.HC) | -1.19 | 0.2366 |
| **Temporal Cortex** |  |  |
| Anterior Insula (L.alns) | -2.92 | 0.0040 |
| Anterior Insula (R.alns) | -3.09 | 0.0024 |
| Posterior Insula (L.plns) | -1.87 | 0.0631 |
| Posterior Insula (R.plns) | -2.01 | 0.0463 |
| Anterior Transverse Temporal Gyrus (L.aTTG*) | -4.77 | 3.90x10<sup>-6</sup> |
| Anterior Transverse Temporal Gyrus (R.aTTG) | -3.15 | 0.0019 |
| Posterior Transverse Temporal Gyrus (R.pTTG) | -2.35 | 0.0198 |
| Posterior Transverse Temporal Gyrus (L.pTTG) | -2.52 | 0.0126 |
| Anterior Superior Temporal Gyrus (L.aSTG) | -2.57 | 0.0111 |
| Anterior Superior Temporal Gyrus (R.aSTG) | -1.53 | 0.1269 |
| Posterior Superior Temporal Gyrus (L.pSTG) | -2.60 | 0.0103 |
| Posterior Superior Temporal Gyrus (R.pSTG) | -3.26 | 0.0014 |
| Anterior Middle Temporal Gyrus (L.aMTG) | -1.74 | 0.0837 |
| Anterior Middle Temporal Gyrus (R.aMTG) | -1.36 | 0.1759 |
| Posterior Middle Temporal Gyrus (L.pMTG) | -1.06 | 0.2888 |
| Posterior Middle Temporal Gyrus (R.pMTG) | -0.95 | 0.3456 |
| Anterior Inferior Temporal Gyrus (L.aITG) | -2.15 | 0.0327 |
| Anterior Inferior Temporal Gyrus (R.aITG) | -2.35 | 0.0201 |
| Posterior Inferior Temporal Gyrus (L.pITG) | -1.83 | 0.0689 |
| Posterior Inferior Temporal Gyrus (R.pITG) | -2.92 | 0.0040 |
| Anterior Fusiform Gyrus (L.aFFG) | -2.12 | 0.0354 |
| Anterior Fusiform Gyrus (R.aFFG) | -2.20 | 0.0291 |
| Posterior Fusiform Gyrus (L.pFFG) | -3.47 | 0.0007 |
| Posterior Fusiform Gyrus (R.pFFG) | -3.49 | 0.0006 |
| **Parietal Cortex** |  |  |
| Superior Postcentral Gyrus (L.sPotCG) | -2.25 | 0.0260 |
| Superior Postcentral Gyrus (R.sPoCG) | -0.50 | 0.6196 |
| Middle Postcentral Gyrus (L.mPoCG) | -1.18 | 0.2409 |
| Middle Postcentral Gyrus (R.mPoCG) | -0.41 | 0.6851 |
| Inferior Postcentral Gyrus (L.iPoCG) | -1.96 | 0.0516 |
| Inferior Postcentral Gyrus (R.iPoCG) | -0.66 | 0.5092 |
| Superior Parietal Lobule (L.SPL) | -2.06 | 0.0412 |
| Superior Parietal Lobule (R.SPL) | -1.25 | 0.2121 |
| Supramarginal Gyrus (L.SMG) | -1.38 | 0.1688 |
| Supramarginal Gyrus (R.SMG) | -1.22 | 0.2222 |
| Angular Gyrus (L.AG) | -2.25 | 0.0257 |
| Angular Gyrus (R.AG) | -1.68 | 0.0942 |
| Parietal Operculum (L.POP) | -1.82 | 0.0699 |
| Parietal Operculum (R.POP) | -0.82 | 0.4149 |
| Paracentral Lobule (L.PCL) | -1.51 | 0.1321 |
| Paracentral Lobule (R.PCL) | -0.63 | 0.5305 |
| Precuneus (L.PCun) | -3.48 | 0.0006 |
| Precuneus (R.PCun)* | -3.93 | 0.0001 |
| **Occipital** |  |  |
| Cuneus (L.Cun) | -1.99 | 0.0485 |
| Cuneus (R.Cun) | -2.66 | 0.0085 |
| Lingual Gyrus (L.LG) | -3.30 | 0.0012 |
| Lingual Gyrus (R.LG)* | -4.11 | 0.0001 |
| Calcarine Sulcus (R.Calc) | -3.04 | 0.0028 |
| Calcarine Sulcus (L.Calc)* | -3.75 | 0.0002 |
| Superior Occipital Gyrus (L.sOG) | -1.85 | 0.0657 |
| Superior Occipital Gyrus (R.sOG) | -2.23 | 0.0267 |
| Middle Occipital Gyrus (L.mOG) | 0.34 | 0.7351 |
| Middle Occipital Gyrus (R.mOG) | -0.75 | 0.4525 |
| Inferior Occipital Gyrus (L.iOG) | -0.53 | 0.5947 |
| Inferior Occipital Gyrus (R.iOG) | -1.19 | 0.2345 |
| **Basal Ganglia** |  |  |
| Caudate Nuclues (L.CN)* | -5.05 | 1.12x10<sup>-6</sup> |
| Caudate Nuclues (R.CN)* | -5.70 | 4.99x10<sup>-8</sup> |
| Nucleus Accumbens (L.NA)* | -4.77 | 3.84x10<sup>-6</sup> |
| Nucleus Accumbens (R.NA)* | -4.23 | 3.81x10<sup>-5</sup> |
| Basal Forebrain Nuclei (L.BF) | -1.25 | 0.2131 |
| Basal Forebrain Nuclei (R.BF) | -1.77 | 0.0792 |
| Putamen (L.Pu)* | -6.18 | 4.55x10<sup>-9</sup> |
| Putamen (R.Pu)* | -6.68 | 3.10x10<sup>-10</sup> |
| Globus Pallidus (L.GP) | 1.68 | 0.0954 |
| Globus Pallidus (R.GP) | 2.72 | 0.0073 |
| Thalamus (L.Tha) | -0.40 | 0.6920 |
| Thalamus (R.Tha) | 1.12 | 0.2653 |
| Hypothalamus (L.HTh) | -1.78 | 0.0762 |
| Hypothalamus (R.HTh) | -1.42 | 0.1570 |
| **Cerebellum** |  |  |
| Cerebellum IX-Tonsil (L.CerIX) | -2.46 | 0.0149 |
| Cerebellum IX-Tonsil (R.CerIX) | -2.83 | 0.0052 |
| Cerebellum VIIIB-Inferior Posterior -PML (L.CerVIIIB) | -2.46 | 0.0147 |
| Cerebellum VIIIB-Inferior Posterior -PML (R.CerVIIIB) | -3.00 | 0.0031 |
| Cerebellum VIIA-Superior Posterior -Crus I (L.CrusI) | -1.29 | 0.1971 |
| Cerebellum VIIA-Superior Posterior -Crus I (R.CrusI) | -1.94 | 0.0543 |
| Cerebellum VIIA-Superior Posterior -Crus II (L.CrusII) | -2.79 | 0.0059 |
| Cerebellum VIIA-Superior Posterior -Crus II (R.CrusII)* | -3.75 | 0.0002 |
| Cerebellum VI-Superior Posterior (L.CerVI) | -3.17 | 0.0018 |
| Cerebellum VI-Superior Posterior (R.CerVI) | -3.38 | 0.0009 |
| Cerebellum V-Anterior B (L.CerVB) | -2.99 | 0.0032 |
| Cerebellum V-Anterior B (R.CerVB) | -3.13 | 0.0020 |
| Cerebellum V-Anterior A (L.CerVA)* | -3.78 | 0.0002 |
| Cerebellum V-Anterior A (R.CerVA)* | -4.91 | 2.09x10<sup>-6</sup> |
| Cerebellum IV-Anterior Quadrangulate (L.CerIV)* | -4.39 | 1.95x10<sup>-5</sup> |
| Cerebellum IV-Anterior Quadrangulate (R.CerIV)* | -5.75 | 4.02x10<sup>-8</sup> |
| Cerebellum III-Anterior Quadrangulate (L.CerIII)* | -3.77 | 0.0002 |
| Cerebellum III-Anterior Quadrangulate (R.CerIII)* | -4.42 | 1.75x10<sup>-5</sup> |
| Cerebellum II-Anterior Quadrangulate (L.CerII) | -3.18 | 0.0018 |
| Cerebellum II-Anterior Quadrangulate (R.CerII) | -2.83 | 0.0052 |

**Key:** L. refers to region in the left hemisphere, while R. the right hemisphere. \* multiple comparisons correction at FWE $p \leq 0.05$ .

**Supplementary Table 3.**
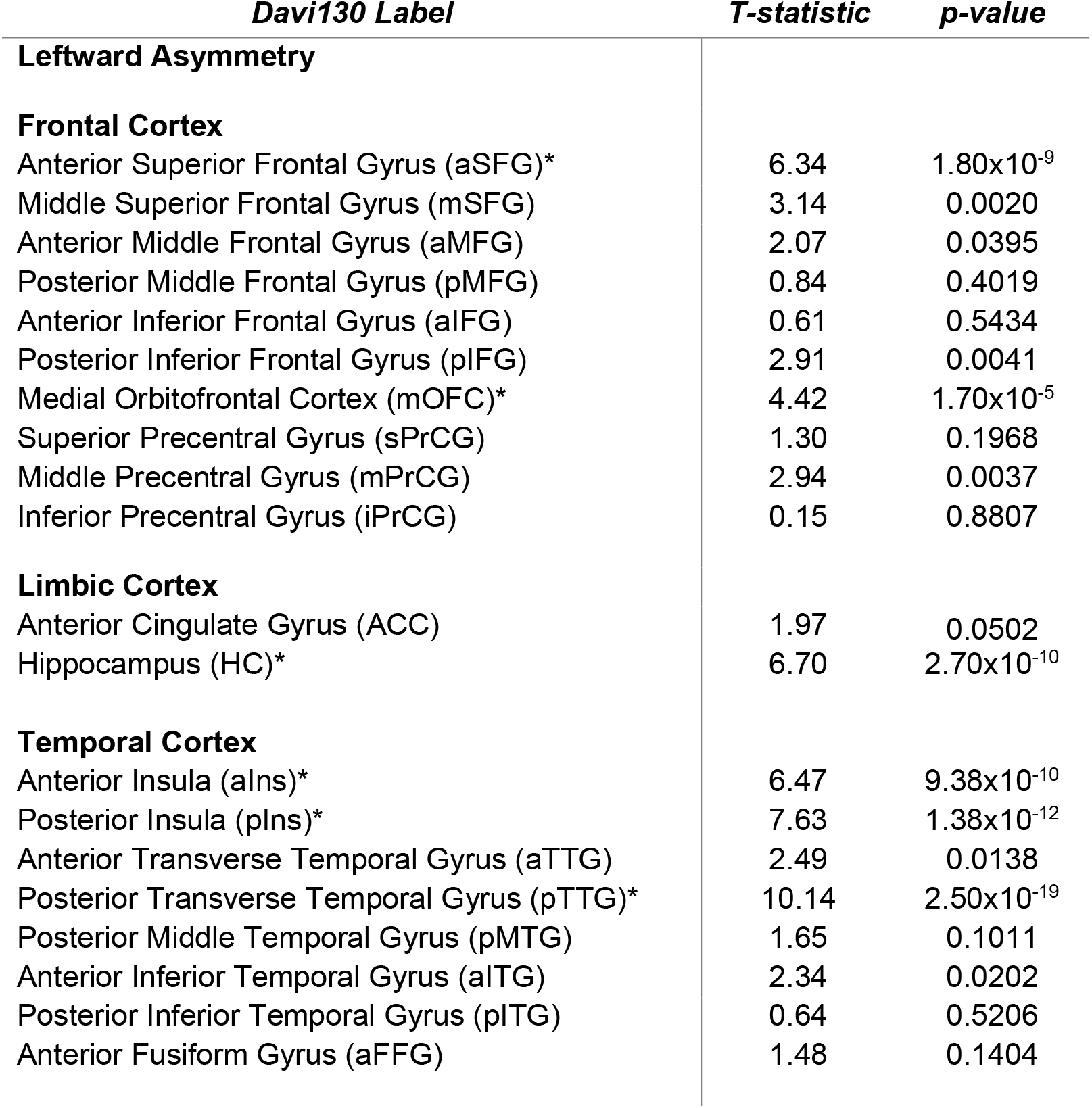
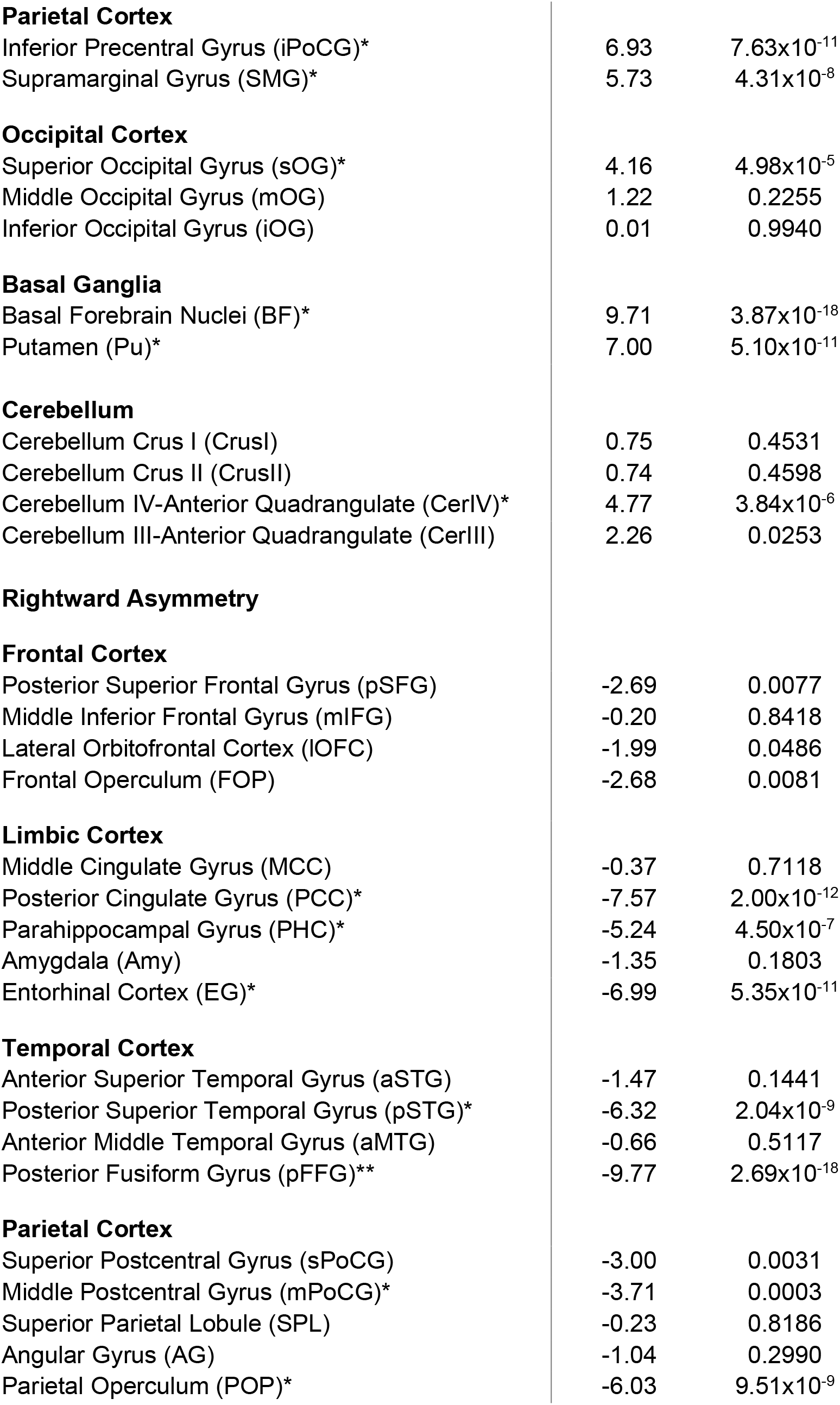
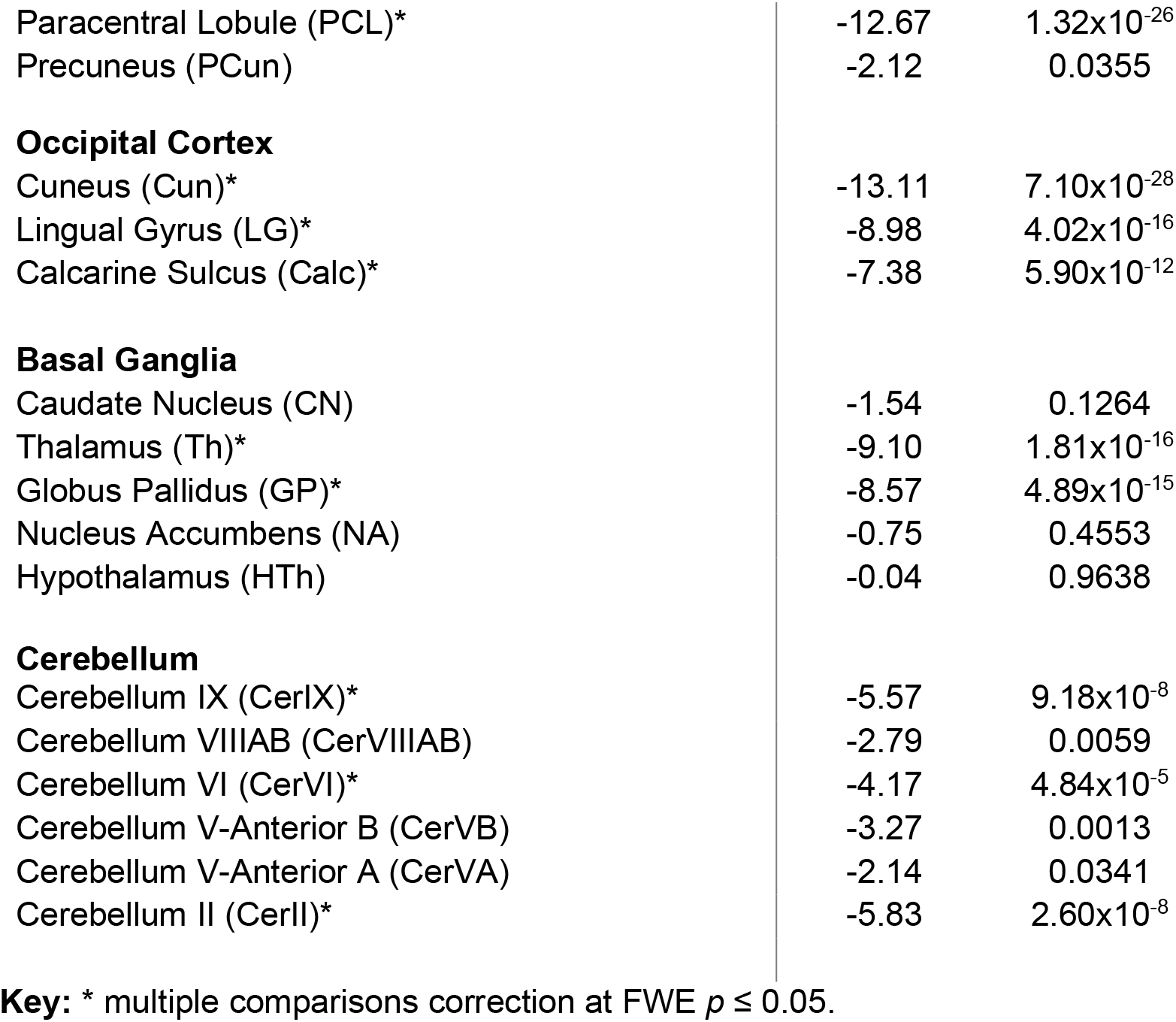
Davi130 labels cortical hemispheric asymmetry

| *Davi130 Label* | *T-statistic* | *p-value* |
| --- | --- | --- |
| **Leftward Asymmetry** |  |  |
| **Frontal Cortex** |  |  |
| Anterior Superior Frontal Gyrus (aSFG)* | 6.34 | 1.80x10<sup>-9</sup> |
| Middle Superior Frontal Gyrus (mSFG) | 3.14 | 0.0020 |
| Anterior Middle Frontal Gyrus (aMFG) | 2.07 | 0.0395 |
| Posterior Middle Frontal Gyrus (pMFG) | 0.84 | 0.4019 |
| Anterior Inferior Frontal Gyrus (aIFG) | 0.61 | 0.5434 |
| Posterior Inferior Frontal Gyrus (pIFG) | 2.91 | 0.0041 |
| Medial Orbitofrontal Cortex (mOFC)* | 4.42 | 1.70x10<sup>-5</sup> |
| Superior Precentral Gyrus (sPrCG) | 1.30 | 0.1968 |
| Middle Precentral Gyrus (mPrCG) | 2.94 | 0.0037 |
| Inferior Precentral Gyrus (iPrCG) | 0.15 | 0.8807 |
| **Limbic Cortex** |  |  |
| Anterior Cingulate Gyrus (ACC) | 1.97 | 0.0502 |
| Hippocampus (HC)* | 6.70 | 2.70x10<sup>-10</sup> |
| **Temporal Cortex** |  |  |
| Anterior Insula (aIns)* | 6.47 | 9.38x10<sup>-10</sup> |
| Posterior Insula (pIns)* | 7.63 | 1.38x10<sup>-12</sup> |
| Anterior Transverse Temporal Gyrus (aTTG) | 2.49 | 0.0138 |
| Posterior Transverse Temporal Gyrus (pTTG)* | 10.14 | 2.50x10<sup>-19</sup> |
| Posterior Middle Temporal Gyrus (pMTG) | 1.65 | 0.1011 |
| Anterior Inferior Temporal Gyrus (aITG) | 2.34 | 0.0202 |
| Posterior Inferior Temporal Gyrus (pITG) | 0.64 | 0.5206 |
| Anterior Fusiform Gyrus (aFFG) | 1.48 | 0.1404 |
| **Parietal Cortex** |  |  |
| Inferior Precentral Gyrus (iPoCG)* | 6.93 | $7.63 \times 10^{-11}$ |
| Supramarginal Gyrus (SMG)* | 5.73 | $4.31 \times 10^{-8}$ |
| **Occipital Cortex** |  |  |
| Superior Occipital Gyrus (sOG)* | 4.16 | $4.98 \times 10^{-5}$ |
| Middle Occipital Gyrus (mOG) | 1.22 | 0.2255 |
| Inferior Occipital Gyrus (iOG) | 0.01 | 0.9940 |
| **Basal Ganglia** |  |  |
| Basal Forebrain Nuclei (BF)* | 9.71 | $3.87 \times 10^{-18}$ |
| Putamen (Pu)* | 7.00 | $5.10 \times 10^{-11}$ |
| **Cerebellum** |  |  |
| Cerebellum Crus I (CrusI) | 0.75 | 0.4531 |
| Cerebellum Crus II (CrusII) | 0.74 | 0.4598 |
| Cerebellum IV-Anterior Quadrangulate (CerIV)* | 4.77 | $3.84 \times 10^{-6}$ |
| Cerebellum III-Anterior Quadrangulate (CerIII) | 2.26 | 0.0253 |
| **Rightward Asymmetry****Frontal Cortex** |  |  |
| Posterior Superior Frontal Gyrus (pSFG) | -2.69 | 0.0077 |
| Middle Inferior Frontal Gyrus (mIFG) | -0.20 | 0.8418 |
| Lateral Orbitofrontal Cortex (IOFC) | -1.99 | 0.0486 |
| Frontal Operculum (FOP) | -2.68 | 0.0081 |
| **Limbic Cortex** |  |  |
| Middle Cingulate Gyrus (MCC) | -0.37 | 0.7118 |
| Posterior Cingulate Gyrus (PCC)* | -7.57 | $2.00 \times 10^{-12}$ |
| Parahippocampal Gyrus (PHC)* | -5.24 | $4.50 \times 10^{-7}$ |
| Amygdala (Amy) | -1.35 | 0.1803 |
| Entorhinal Cortex (EG)* | -6.99 | $5.35 \times 10^{-11}$ |
| **Temporal Cortex** |  |  |
| Anterior Superior Temporal Gyrus (aSTG) | -1.47 | 0.1441 |
| Posterior Superior Temporal Gyrus (pSTG)* | -6.32 | $2.04 \times 10^{-9}$ |
| Anterior Middle Temporal Gyrus (aMTG) | -0.66 | 0.5117 |
| Posterior Fusiform Gyrus (pFFG)** | -9.77 | $2.69 \times 10^{-18}$ |
| **Parietal Cortex** |  |  |
| Superior Postcentral Gyrus (sPoCG) | -3.00 | 0.0031 |
| Middle Postcentral Gyrus (mPoCG)* | -3.71 | 0.0003 |
| Superior Parietal Lobule (SPL) | -0.23 | 0.8186 |
| Angular Gyrus (AG) | -1.04 | 0.2990 |
| Parietal Operculum (POP)* | -6.03 | $9.51 \times 10^{-9}$ |
| Paracentral Lobule (PCL)* | -12.67 | 1.32x10<sup>-26</sup> |
| Precuneus (PCun) | -2.12 | 0.0355 |
| **Occipital Cortex** |  |  |
| Cuneus (Cun)* | -13.11 | 7.10x10<sup>-28</sup> |
| Lingual Gyrus (LG)* | -8.98 | 4.02x10<sup>-16</sup> |
| Calcarine Sulcus (Calc)* | -7.38 | 5.90x10<sup>-12</sup> |
| **Basal Ganglia** |  |  |
| Caudate Nucleus (CN) | -1.54 | 0.1264 |
| Thalamus (Th)* | -9.10 | 1.81x10<sup>-16</sup> |
| Globus Pallidus (GP)* | -8.57 | 4.89x10<sup>-15</sup> |
| Nucleus Accumbens (NA) | -0.75 | 0.4553 |
| Hypothalamus (HTh) | -0.04 | 0.9638 |
| **Cerebellum** |  |  |
| Cerebellum IX (CerIX)* | -5.57 | 9.18x10<sup>-8</sup> |
| Cerebellum VIIIAB (CerVIIIAB) | -2.79 | 0.0059 |
| Cerebellum VI (CerVI)* | -4.17 | 4.84x10<sup>-5</sup> |
| Cerebellum V-Anterior B (CerVB) | -3.27 | 0.0013 |
| Cerebellum V-Anterior A (CerVA) | -2.14 | 0.0341 |
| Cerebellum II (CerII)* | -5.83 | 2.60x10<sup>-8</sup> |

**Key:** \* multiple comparisons correction at FWE $p \leq 0.05$ .

